# Aspects of the ecology of *Bartramia halleriana* and *Crossocalyx hellerianus* in oceanic deciduous woodland

**DOI:** 10.1101/2020.04.28.066522

**Authors:** Des A. Callaghan, Jamie Bevan

**Author notes:** Correspondence to: Des A. Callaghan, 20 Thornleigh Road, Bristol, BS7 8PH, UK.

## Abstract

**Introduction:** This study investigates the ecology of two boreal-montane bryophytes, the saxicolous moss *Bartramia halleriana* and the epixylic liverwort *Crossocalyx hellerianus*, at the edge of their ranges, in oceanic deciduous woodland.

**Methods:** The study site comprises two adjoining woodlands, Allt Penyrhiw-iar (canopy trees ca. 100 yr old) and Allt Rhyd Groes (ca. 200 yr), Carmarthenshire, UK. The distribution and abundance of *B. halleriana* and *C. hellerianus* was surveyed, relevés sampled to record habitat and community composition, and sporophyte frequency and stage of development measured. Light climate of *B. halleriana* was investigated via hemispherical photography, and abundance of large rotten logs used as a measure of habitat quality for *C. hellerianus*.

**Results and discussion:** Four subpopulations of *B. halleriana* occur, comprising 21 individual-equivalents (occupied 1 m grid cells), all on mildly base-rich mudstone of north-facing rockfaces, with very little direct solar radiation and a diverse assemblage of bryophytes. Sporophyte are scarce. A total of 143 individual-equivalents (occupied logs or trees) of *C. hellerianus* was recorded, an exceptional population in Wales. Most of the population is within Allt Rhyd Groes, where large rotten logs are much more abundant due to greater woodland age. The liverwort mainly occupied the sides of decorticated rotten logs, amongst a sparse community composed mainly of other small liverworts and *Cladonia* lichens. Neither sporophytes nor perianths were found. Further research could usefully focus on the description and measurement of favourable habitat for each species and associated physiological tolerances.

## Introduction

Sites that are rich in rare and scarce species are of special conservation value and there is a clear need to ensure their protection and effective management. Indeed, criteria used to select sites for safe-guarding commonly include reference to such species, for example regarding ‘restricted-range species’ in the context of ‘key biodiversity areas’ (Eken et al. 2004). In Britain, the statutory nature conservation agencies have a duty under the Wildlife and Countryside Act 1981 (as amended), to notify any area of land which in their opinion is ‘of special interest by reason of any of its flora, fauna, or geological or physiographical features’ (Bainbridge et al. 2013). These areas are known as Sites of Special Scientific Interest (SSSIs). Some of the criteria used for the identification of SSSIs are based on a scoring system, with thresholds to determine occurrence within sites of nationally important assemblages of rare and scarce species, including bryophytes (Bosanquet et al. 2018). Due to their national scarcity (Pescott 2016), *Bartramia halleriana* Hedw. (Bartramiales: Bartramiaceae) and *Crossocalyx hellerianus* (Nees ex Lindenb.) Meyl (Jungermanniales: Anastrophyllaceae) can contribute to scores used for the identification of SSSIs in Britain, and as a consequence they can become a focus of site-based monitoring and management activities, for which knowledge of their ecology is fundamental. The monoicous moss *B. halleriana* is a boreal-montane species (Hill and Preston 1998), occurring frequently in some parts of Europe and Asia, and very rarely in North America, Central America and Africa (Blockeel et al. 2014; Fransén 2004). In Britain, it was recorded in 57 10 km grid cells during 1990–2013, in northern England, Scotland and Wales (Blockeel et al. 2014). The dioicous liverwort *C. hellerianus* also exhibits a boreal-montane distribution pattern (Hill and Preston 1998), essentially circumboreal and largely continental, occurring in Europe, Asia and North America (Blockeel et al. 2014; Hong 1996; Sabovljevic et al. 2019; Schuster 1969; Schill and Long 2003). Its range in Britain is similar to *B. halleriana*, being recorded in 38 10 km grid cells during 1990–2013, confined to northern England, Scotland and Wales (Blockeel et al. 2014). To date, no research has been undertaken regarding the ecology of either species in Britain. The purpose of this study is to investigate such, to help inform conservation decisions and research priorities.

## Method

### Study area

The study site (52°07’05”N, 3°48’37”W; 160–350 m a.s.l.) comprises two contiguous woodland compartments, Allt Penyrhiw-iar (17.6 ha) and Allt Rhyd Groes (14.6 ha), Carmarthenshire (v.-c. 44), South Wales. It supports a nationally important assemblage of rare and scarce bryophytes of woodland, including *B. halleriana* and *C. hellerianus*. The site is within Allt Rhyd y Groes National Nature Reserve (NNR), and the much larger protected areas of Cwm Doethie–Mynydd Mallaen SSSI and Cwm Doethie–Mynydd Mallaen Special Area of Conservation (SAC). The woodland mainly comprises W17 *Quercus petraea–Betula pubescens–Dicranum majus* woodland and W11 *Quercus petraea–Betula pubescens–Oxalis acetosella* woodland (Rodwell 1991), located on steep hillsides, with geology dominated by mudstones, weathering to produce clay soils. The climate is oceanic, with 176 rain-days yr^-1^ (days with >1 mm rain) and average temperatures of 14.6°C during the hottest month (July) and 3.3°C during the coldest month (February) for the period 1961–2002 (Met Office data supplied through the UK Climate Impact Programme). The NNR was established in 1959 and has since been managed for nature conservation, with a general policy of low intervention woodland management. Conservation management has recently included thinning of understorey trees and shrubs in some parts and controlled light grazing by sheep (ca. 0.05 LSU ha^-1^ yr^-1^) is practised throughout. Little is known of the long-term management history of the two woodland compartments within the study area, though in 1821, 249 oaks from ‘Allt Rhyd y Groes’ were sold by Cawdor Estate, the landowner at that time, possibly for ship building (NRW 2020). Indeed, canopy oak trees in both compartments are relatively even-aged, indicating clear-felling in the past, perhaps followed by re-planting. However, those in Allt Rhyd Groes average ca. 200 yr old, while those in Allt Penrhiw-iar are mostly <100 yr (NRW 2020). Both woodlands have been identified as ‘ancient woodland’ (Lister and Whitebread 1988) and are likely to have a long history of tree cover, notwithstanding episodes of felling for timber.

### Taxonomy

Taxonomy follows Hodgetts et al. (2020) for bryophytes, Smith et al. (2009) for lichens and Stace (2010) for vascular plants.

### Geographic reference system

Geographic coordinates follow the Ordnance Survey (OS) National Grid reference system (EPSG: 27700). Grid cells are referred to by the coordinates of the south-west corner.

### Distribution and abundance

COSWIC (2011) defined an individual of *B. halleriana* as “a discrete colony (clump or tuft of moss consisting of many shoots)”. Following trial in the field, including further definition of “discrete” (≥10 cm distance from any other count unit) and “many shoots” (≥10), this was not followed presently because it ignored frequent occurrences of the moss that did not fit within the definition. Indeed, whilst *B. halleriana* is a moss with a ‘tuft’ growth form (Hill et al. 2007), colonies frequently have fuzzy boundaries, for example with tufts sometimes blending into one another, larger tufts sometimes becoming fragmented due to disturbance events, and other occurrences not being tufts at all, but rather scattered shoots amongst other mosses, making the counting of discrete colonies troublesome. Instead, an ‘individual-equivalent’ of *B. halleriana* was here defined as an occupied 1 m grid cell, following Bergamini et al. (2019). Such were counted after marking-out locations of the moss in the field, and where necessary using a tape-measure to determine number of occupied 1 m grid cells. An individual-equivalent of *C. hellerianus* is defined as an occupied log or tree, again following Bergamini et al. (2019). An inventory of previously known locations of both species within the study area was compiled from records within the national recording database of the British Bryological Society (BBS), held by the Biological Records Centre (Wallingford, UK), and reports of previous bryophyte surveys (Newton 1997, 2002, 2008). During September 2019 and February–March 2020, the study site was searched for locations of the two species, revisiting all former locations and ensuring good coverage of other areas. The survey trail was logged with a hand-held GPS unit (Garmin GPSMAP 64s, Garmin Ltd, Olathe, USA), which generally reported an accuracy of ≤6 m during the survey conditions. When an individual-equivalent of either species was found, coordinates of the location were logged.

### Habitat and community composition

During 26 February to 3 March 2020, relevés (50 × 25 cm) were recorded for *C. hellerianus* (*n* = 10) and *B. halleriana* (*n* = 10), to describe habitat conditions and community composition, generally following the method of Bates (2011). Sample locations were chosen to represent the full range of conditions occupied by each species. Percentage cover of each species of bryophyte, vascular plants, lichen and macroalgae was estimated, as was the percentage cover of dead plant material (‘litter’) and bare ground. Shade was recorded according to the following index: 1, fully exposed to sunlight at all times; 2, shaded from direct sunlight for up to half the day; 3, receiving significant direct sunlight but for less than half the day; 4, moderately shaded from direct sunlight, e.g. by a light-medium deciduous tree canopy; 5, permanently shaded from direct sunlight but otherwise open to the sky (i.e. with north-facing aspect); 6, in deep woodland (e.g. coniferous) shade with no sunflecks; 7, in perpetual, very deep shade. Slope was measured with a digital clinometer, recorded as the average angle (°) from horizontal in the direction of greatest slope. Aspect was recorded as the bearing (°) of the relevé in the direction of greatest slope using the above GPS unit.

### Sporophyte frequency and stage of development

Within the above relevés, a careful search was undertaken for sporophytes of *B. halleriana*, and counts made of any found, categorised as: A – calyptra partially exerted; B – capsule beginning to expand; C – green capsule widened fully, lid intact; D – capsule brown, lid intact; E – capsule brown, lid detached; F – capsule predated; or G – sporophyte from previous season. For *C. hellerianus*, a careful search was made for perianths and sporophytes.

### Bartramia halleriana *light climate*

At a representative colony of *B. halleriana*, a 180° hemispherical sky view image was captured with a Laowa 4mm f2.8 fisheye lens mounted on an Olympus E-M10ii camera body. The camera was levelled using inbuilt gauges, and the bearing (°) of the image recorded with a digital compass. The image was processed in Adobe Photoshop, including its rotation to align magnetic north with the top of the image, and addition of sun tracks, paths of which were derived from ‘SunEarthTools’ (www.sunearthtools.com).

### Crossocalyx hellerianus *habitat quality*

To investigate habitat quality for *C. hellerianus* in Allt Penyrhiw-iar vs. Allt Rhyd Groes, ten random plots, each an OS 50 m grid cell, were sampled in each woodland. Each grid cell was visited, and a count made of the number of large rotten logs. Logs were counted only if they were lying on the ground, decorticated and in an advanced stage of decay, and dimensions exceeded 25 cm diameter at base and 5 m length. Such logs overlapping the boundary of a grid cell were counted if crossing the north or west boundary; those crossing the south or east boundary were ignored. The resultant data deviated significantly from a normal distribution, indicated by a Shapiro–Wilk test, and so a Mann-Whitney-Wilcoxon test was used to test for significant difference in counts of logs in Allt Penyrhiw-iar vs. Allt Rhyd Groes.

## Results

### Distribution and abundance

Survey coverage in each woodland compartment was similar, totalling 14.1 km (0.80 km ha^-1^) in Allt Penyrhiw-iar and 12.5 km (0.86 km ha^-1^) in Allt Rhyd Groes. Four discrete subpopulations of *B. halleriana* were found (Figure 1A), comprising 21 individual-equivalents, most (90%; *n* = 19) within two subpopulations (Figure 2). A total of 143 individual-equivalents of *C. hellerianus* was recorded, far more in Allt Rhyd Groes (10 individual-equivalents km^-1^; *n* = 125; 87%) compared to Allt Penyrhiw-iar (1.3 individual-equivalents km^-1^; *n* = 18; 13%) (Figure 1B).

**Figure 1.**
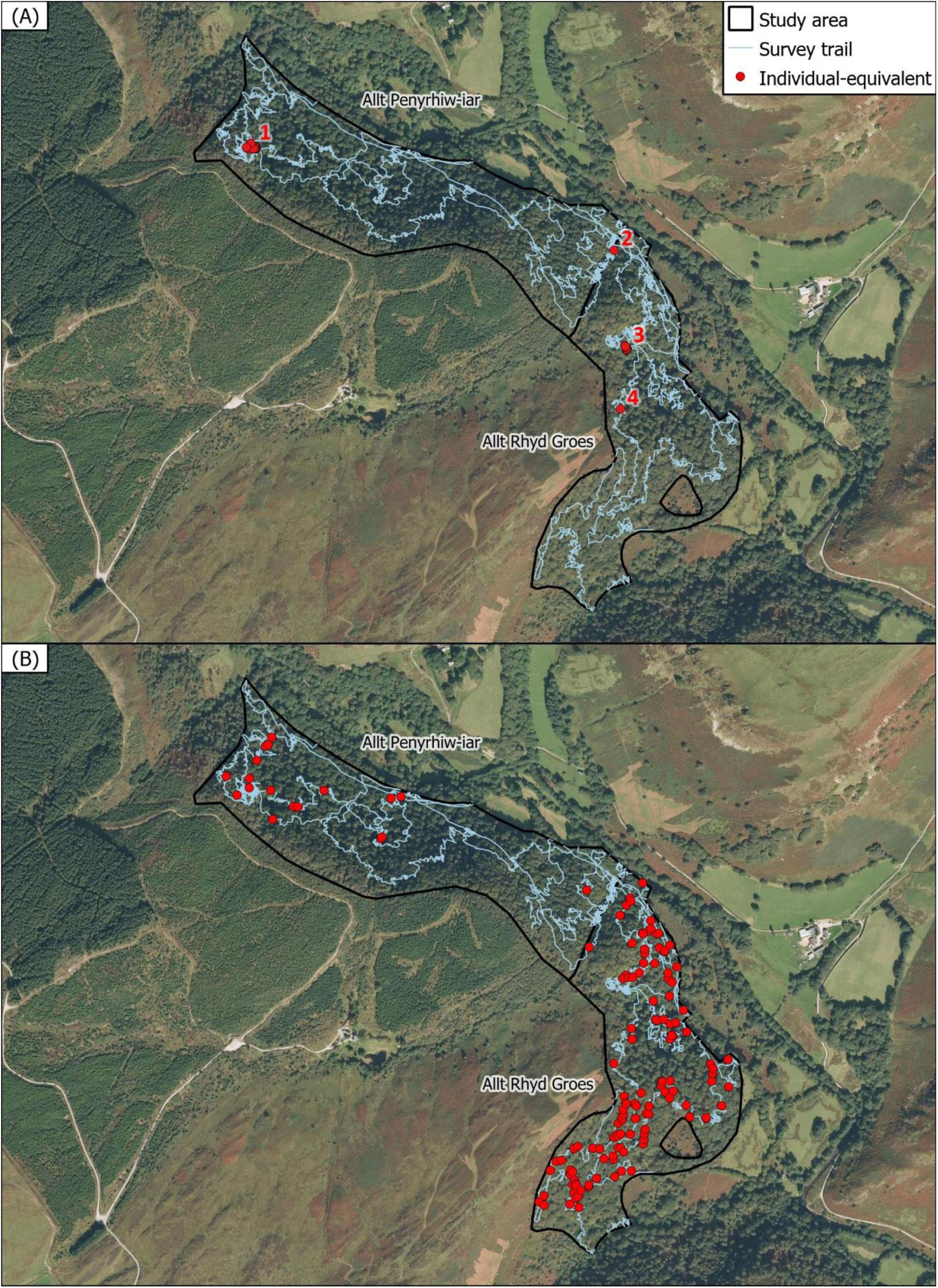
Locations of (A) *Bartramia halleriana*, with numbered subpopulations, and (B) *Crossocalyx hellerianus* within the study site. Satellite imagery ©2020 NASA, TerraMetrics.

**Figure 2.**
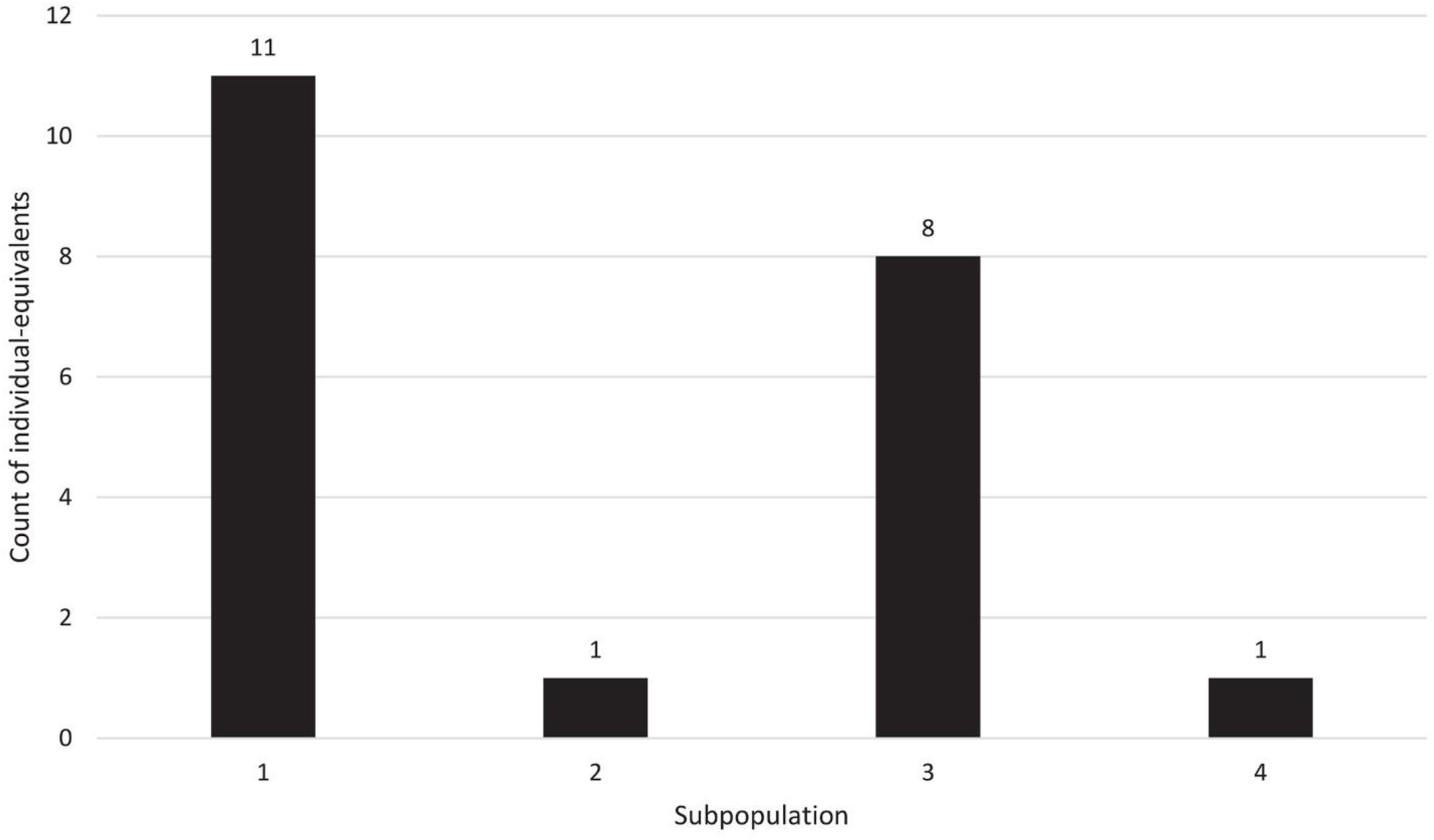
Counts of individual-equivalents (occupied 1 m grid cells) of *Bartramia halleriana* within each subpopulation of the study site.

### Habitat and community composition

Figure 3 shows typical habitat occupied by *B. halleriana* within the study site, and Table 1 shows the percentage cover of species and associated data from relevés (*n* = 10). Bare ground is scarce and the community is diverse, with 40 species recorded in relevés, mostly mosses (*n* = 27; 68%). No macroalgae were present, and both lichens and vascular plants are uncommon. The most frequent associates are the liverworts *Diplophyllum albicans* and *Saccogyna viticulosa*, and the mosses *Heterocladium heteropterum* and *Mnium hornum*. Figure 4 is a polar plot of aspect and slope of relevés, showing confinement of *B. halleriana* within the study area to vertical rock faces (mean slope = 86°; range = 74–95°; *n* = 10) with a northerly aspect.

**Table 1.**
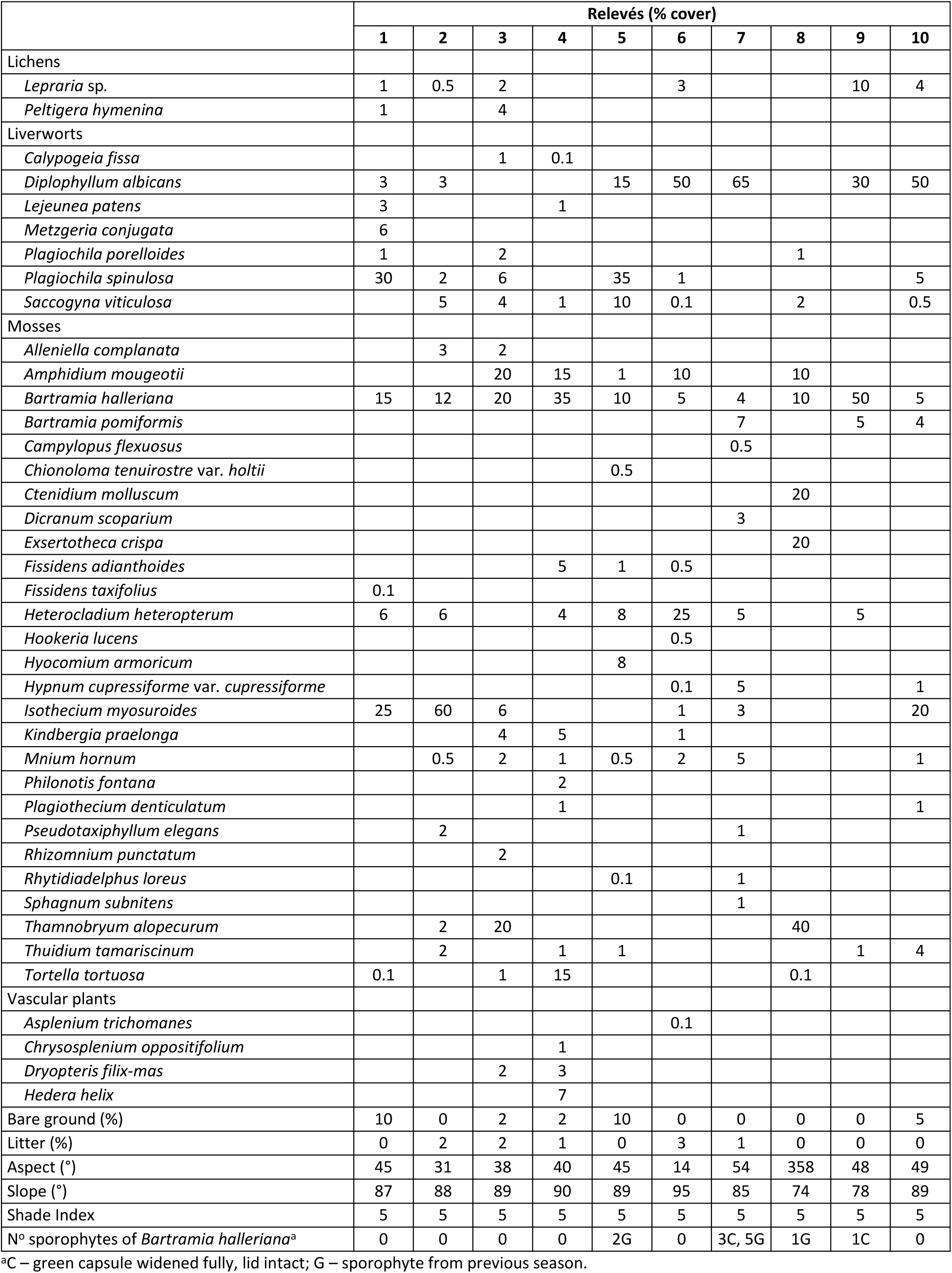
Percentage cover of plant species and associated data from relevés occupied by *Bartramia halleriana* (*n* = 10) within the study area, sampled during 26 February to 3 March 2020. No macroalgae were present.

**Figure 3.**
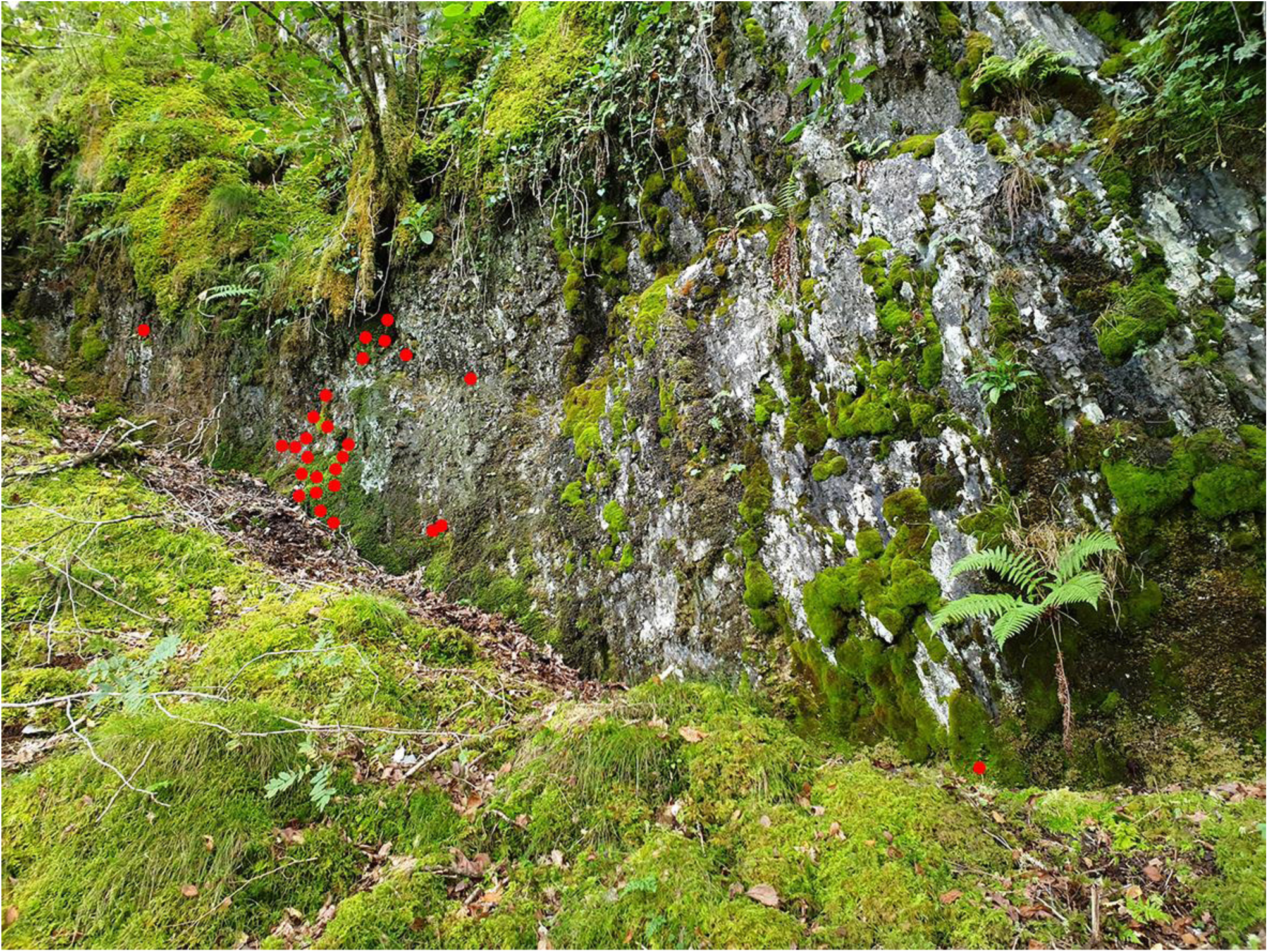
Typical habitat of *Bartramia halleriana* (red dots) within the study site, SN7595148224, 19 September 2019. Photo: D.A. Callaghan.

**Figure 4.**
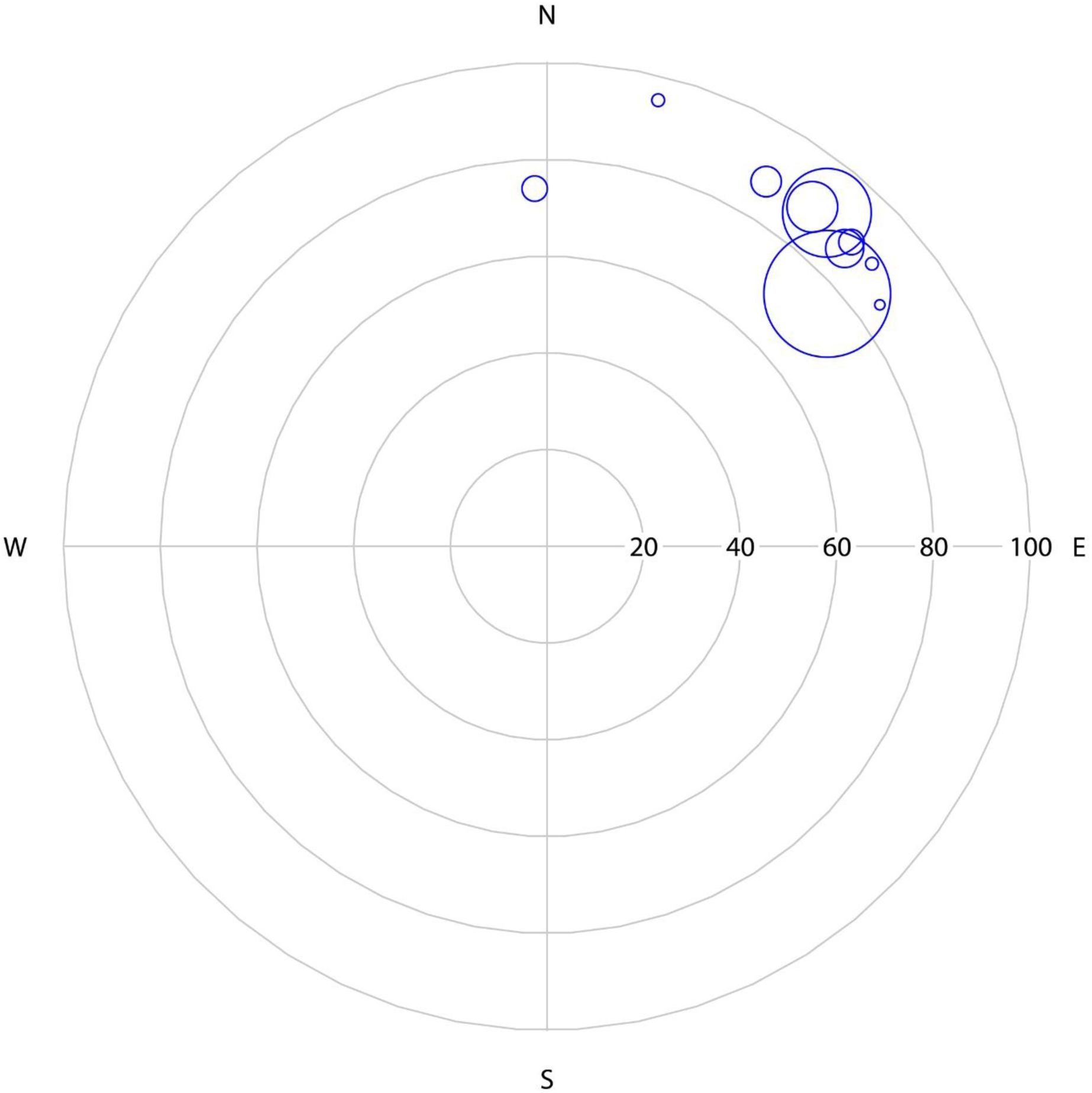
Polar plot illustrating aspect and slope of ten relevés occupied by *Bartramia halleriana*. Dots are scaled by percentage cover of *B. halleriana*.

Figure 5 shows frequency of occurrence of *C. hellerianus* across habitat types occupied within the study site. Rotten logs on the ground support most of the population (*n* = 104; 73%), trunks of *Quercus petraea* are occupied occasionally (*n* = 30; 21%), and standing deadwood (*n* = 5; 3.5%) and trunks of *Betula pubescens* (*n* = 4; 2.8%) are occupied rarely. Figure 6 illustrates the two main types of habitat occupied, and Table 2 shows the percentage cover of species and associated data from relevés on rotten logs (*n* = 10). The rotten log community is sparse, with large areas of bare substrate (mean cover = 70%, range = 50–80%), and species poor, with 16 species recorded in relevés, frequently appearing mottled grey due to squamules of *Cladonia* species (Figure 6A). The most common associates are *Cladonia squamosa, Lepidozia reptans, Nowellia curvifolia* and *Odontoschisma denudatum*. No macroalgae were present, and vascular plants are virtually absent. Small amounts of mosses occur regularly, most often *Campylopus flexuosus*. Figure 7 is a polar plot of aspect and slope of relevés on rotten logs, showing no aspect preference, but illustrating typical occurrence on the sides and over-hanging surfaces of logs, being absent from the tops of logs, where *N. curvifolia* is usually dominant.

**Table 2.** Percentage cover of plant species and associated data from relevés occupied by *Crossocalyx hellerianus* on rotten logs (*n* = 10) within the study area, sampled during 26 February to 3 March 2020. No macroalgae were present.

|  | Relevés (% cover) |  |  |  |  |  |  |  |  |  |
| --- | --- | --- | --- | --- | --- | --- | --- | --- | --- | --- |
|  | 1 | 2 | 3 | 4 | 5 | 6 | 7 | 8 | 9 | 10 |
| Lichens |  |  |  |  |  |  |  |  |  |  |
| *Cladonia coniocraea* |  |  |  |  | 1 |  |  |  |  |  |
| *Cladonia floerkeana* |  |  |  | 0.5 |  |  |  |  |  |  |
| *Cladonia polydactyla* | 2 | 2 |  | 1 | 3 |  |  |  | 3 |  |
| *Cladonia squamosa* | 8 | 5 | 8 | 10 | 10 | 8 | 20 | 3 | 10 | 10 |
| Liverworts |  |  |  |  |  |  |  |  |  |  |
| *Crossocalyx hellerianus* | 1 | 5 | 3 | 5 | 3 | 3 | 2 | 3 | 2 | 3 |
| *Lepidozia reptans* | 1 | 2 | 1 | 2 | 5 | 5 |  | 2 | 3 | 1 |
| *Nowellia curvifolia* | 5 | 5 | 10 | 10 | 20 | 3 | 3 | 10 | 10 |  |
| *Odontoschisma denudatum* | 5 | 2 |  | 5 | 2 | 2 | 0.1 |  | 4 | 5 |
| *Scapania gracilis* |  |  |  | 3 |  |  |  | 2 |  |  |
| *Tritomaria exsectiformis* |  |  |  |  |  |  |  |  | 0.1 |  |
| Mosses |  |  |  |  |  |  |  |  |  |  |
| *Campylopus flexuosus* | 0.1 | 1 | 5 | 5 |  | 0.1 |  | 0.5 | 4 |  |
| *Dicranum scoparium* |  | 0.1 | 3 | 2 |  |  | 1 |  |  | 1 |
| *Hypnum cupressiforme* var. *cupressiforme* |  |  | 1 |  | 1 |  |  |  |  |  |
| *Orthodontium lineare* |  |  |  | 0.1 |  |  |  |  |  |  |
| *Tetraphis pellucida* |  | 3 | 1 |  | 5 |  |  |  |  | 1 |
| Vascular plants |  |  |  |  |  |  |  |  |  |  |
| *Festuca ovina* |  |  | 1 |  |  |  |  |  |  |  |
| Bare substrate (%) | 78 | 75 | 67 | 57 | 50 | 79 | 74 | 80 | 64 | 79 |
| Litter (%) | 0 | 0 | 0 | 0 | 0 | 0 | 0 | 0 | 0 | 0 |
| Aspect (°) | 286 | 261 | 320 | 312 | 47 | 247 | 357 | 36 | 198 | 178 |
| Slope (°) | 64 | 79 | 62 | 100 | 109 | 83 | 107 | 84 | 103 | 82 |
| Shade Index | 4 | 4 | 4 | 4 | 4 | 4 | 4 | 4 | 4 | 4 |
| Log diameter at relevé (cm) | 14 | 18 | 27 | 30 | 32 | 30 | 22 | 26 | 20 | 30 |
| N° perianths of *Crossocalyx hellerianus* | 0 | 0 | 0 | 0 | 0 | 0 | 0 | 0 | 0 | 0 |
| N° sporophytes of *Crossocalyx hellerianus* | 0 | 0 | 0 | 0 | 0 | 0 | 0 | 0 | 0 | 0 |

**Figure 5.**
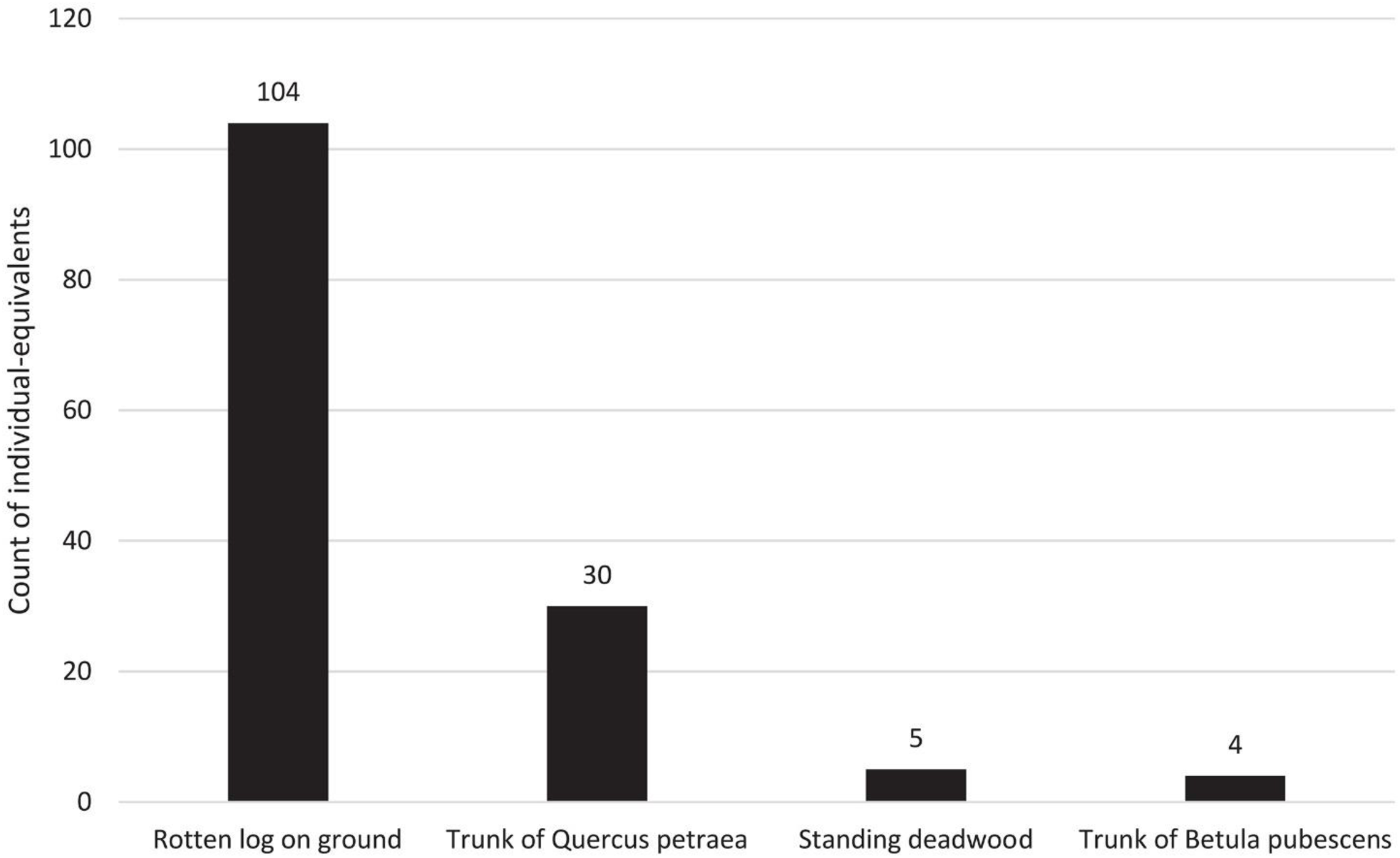
Frequency of occurrence of individual-equivalents (occupied logs or trees) of *Crossocalyx hellerianus* across habitat types within the study site.

**Figure 6.**
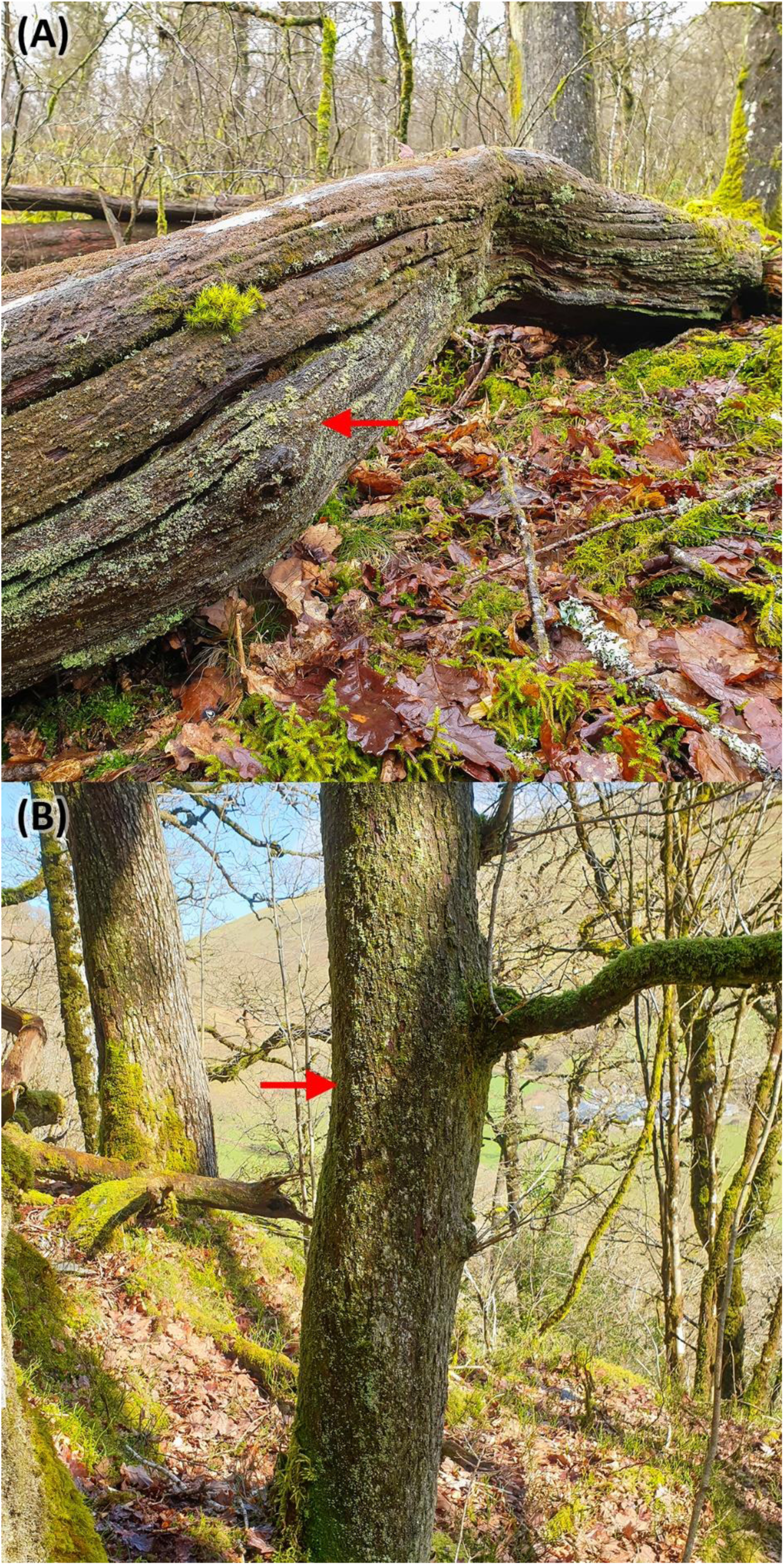
Typical habitat occupied by *Crossocalyx hellerianus* (red arrows) within the study site, including (A) rotten log on ground (SN7677947701, 27 February 2020) and (B) trunk of *Quercus petraea* (SN7672947855, 27 February 2020). Photo: D.A. Callaghan.

**Figure 7.**
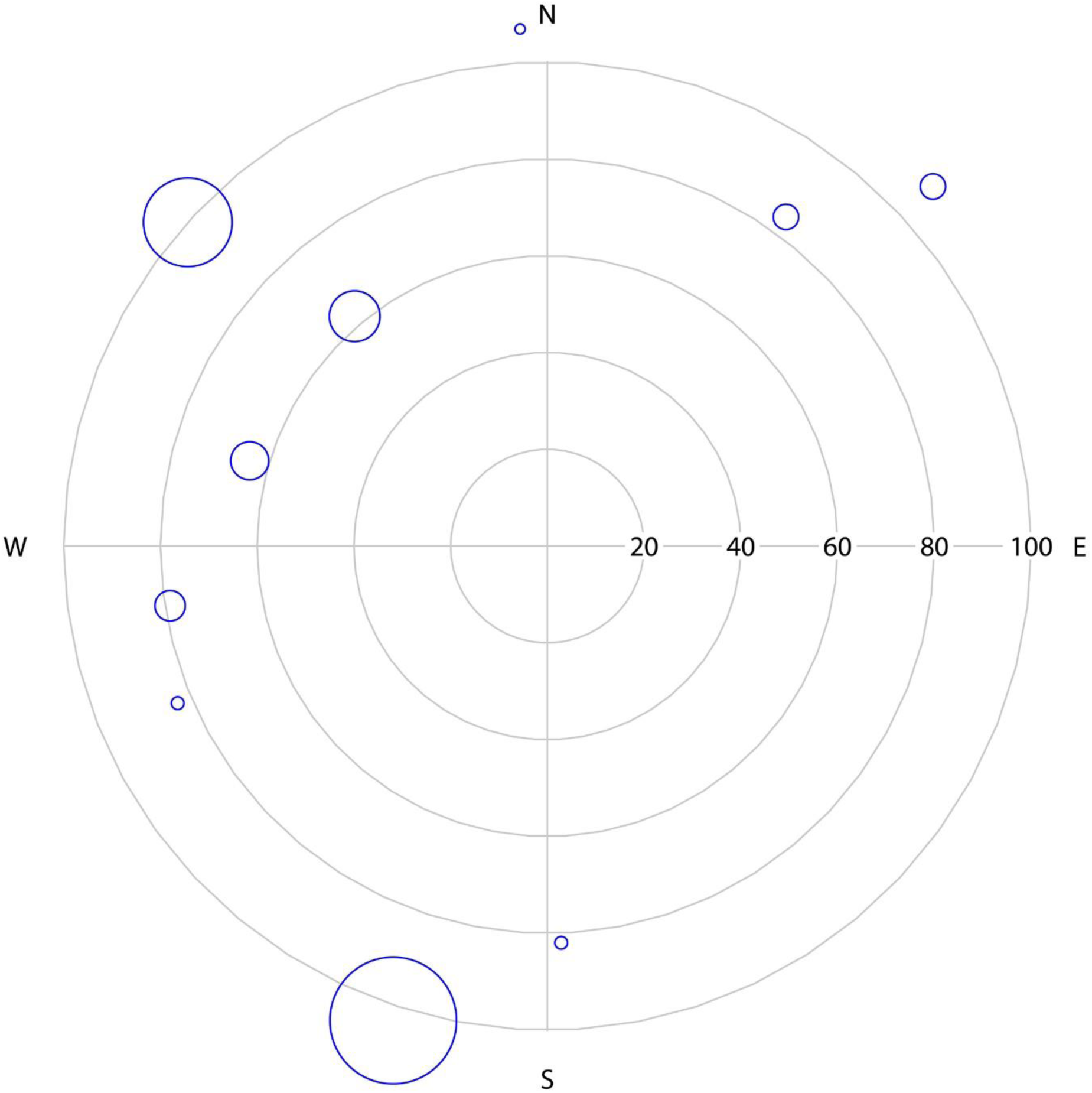
Polar plot illustrating aspect and slope of ten relevés occupied by *Crossocalyx hellerianus* on rotten logs. Dots are scaled by percentage cover of *C. hellerianus*.

### Sporophyte frequency and stage of development

Sporophytes of *B. hellerianum* were scarce, with most relevés lacking any (*n* = 6; 60%). A total of twelve was found in the other relevés, including only four from the current growth season. The latter all had capsules fully grown, but still green and with lids intact. No perianths or sporophytes of *C. hellerianus* were found, either within relevés or elsewhere.

### Bartramia halleriana *light climate*

Figure 8 shows a hemispherical sky view image at a representative colony of *Bartramia halleriana*, with solar tracks overlaid. It shows that during September–March, no direct sunlight reaches this colony, and during April-August, when trees are in-leaf, such events will be limited to sunflecks occurring only for a brief period in early morning.

**Figure 8.**
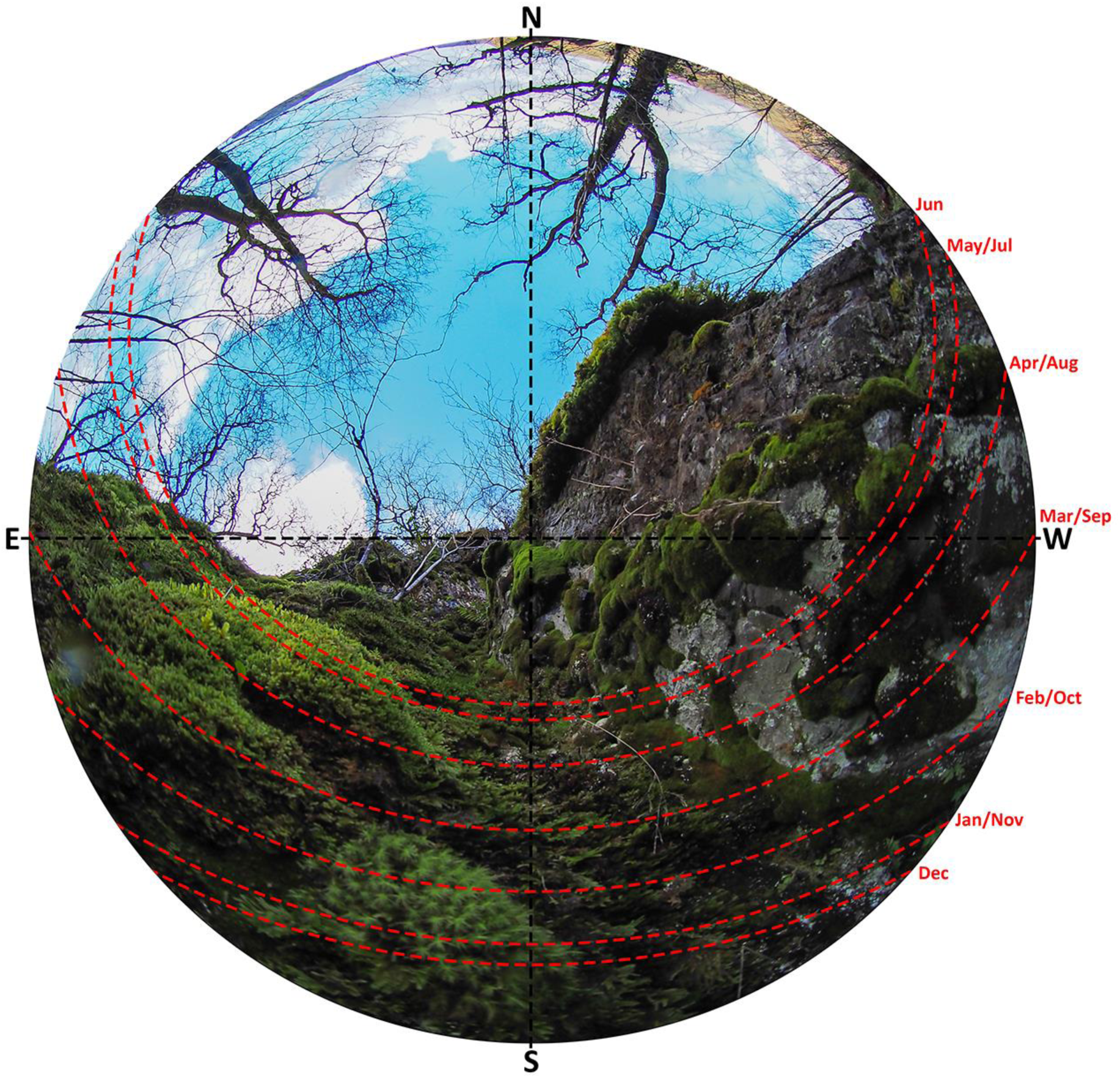
Hemispherical sky view image at a representative colony of *Bartramia halleriana* (red arrow), 9 March 2020, SN7594648227. The margin of the image represents the horizon and the centre is the zenith. Solar tracks are for the 22nd day of each month. Photo: D.A. Callaghan.

### Crossocalyx hellerianus *habitat quality*

Data from random plots show large rotten logs are significantly more abundant in Allt Rhyd Groes compared to Allt Penyrhiw-iar, averaging 11.2 plot^-1^ (44.8 ha^-1^) vs. 1.3 plot^-1^ (5.2 ha^-1^) (*W* = 5, *p* < 0.01) (Figure 9).

**Figure 9.**
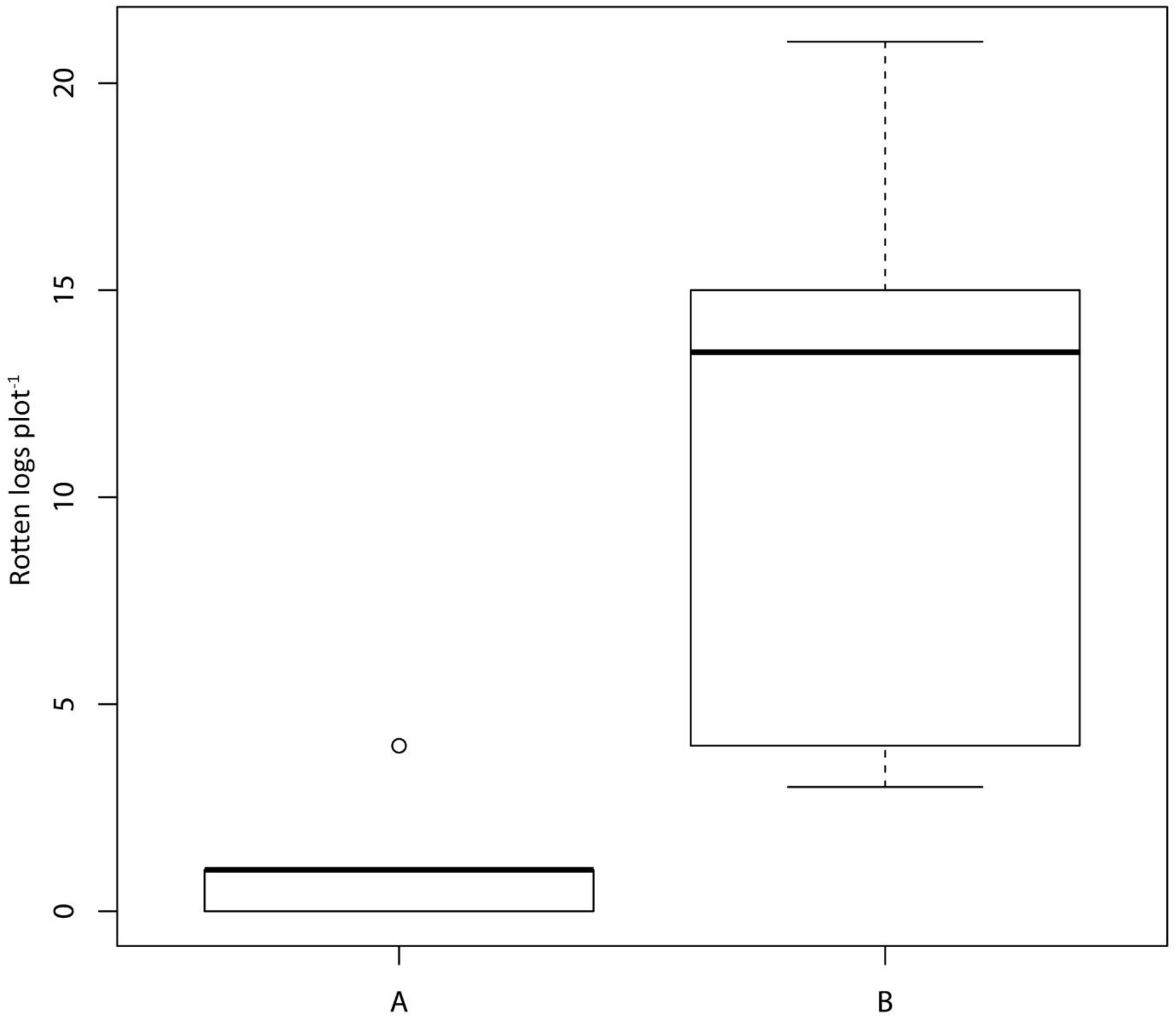
Boxplot of counts of large rotten logs in random OS 50 × 50 m plots (*n* = 20) in (A) Allt Penyrhiw-iar and (B) Allt Rhyd Groes.

## Discussion

*Bartramia halleriana* has been known from the study site since 1906 and results of the present study confirm its continued presence, albeit limited to four small subpopulations, totalling 21 occupied 1 m grid cells. No surveys in Britain have yet documented a larger population, though unfortunately census information is extremely limited. Gwenffrwd Gorge, located 2 km south-west of the present study site, was considered the best site for the species in Carmarthenshire by Bosanquet and Motley (2004), supporting at least four subpopulations, though their size has not been measured. Newton (1997, 2002, 2008) located two of the subpopulations in the study site and it appears that, at least over recent decades, the overall site population has remained reasonably stable. Detailed surveys in Canada, where it is a rare species of conservation concern, report populations that are generally small, but one that totals 284 discrete colonies (Achuff et al. 2010; COSEWIC 2011), far larger than any censused in Britain. The rock occupied by *B. halleriana* within the study area is mudstone and is mildly base-rich, as indicated by occasional presence in relevés of calcicoles, such as *Alleniella complanata, Ctenidium molluscum, Exsertotheca crispa, Thamnobryum alopecurum* and *Tortella tortuosa*. Some calcareous content is typical of rock inhabited by the species in Britain (Blockeel et al. 2014; Hill et al. 2007), Fennoscandia (Hallingbäck et al. 2008) and Ireland (Lockhart et al. 2012), though in Canada it is restricted to non-calcareous rock (COSWIC 2011) and is known from acidic rock elsewhere (Fransén 2004). All colonies within the study area occupy north-facing rock outcrops, where direct solar radiation is very limited. A humid and shaded niche is typical of the species generally, usually beneath a woodland canopy, either coniferous or deciduous (COSWIC 2011; Fransén 2004). This suggests *B. halleriana* may be shade-adapted, perhaps with photoinhibition occurring at relatively low light levels, and possesses limited desiccation tolerance, which deserve investigation.

No phenological studies of *B. halleriana* appear to have been undertaken. This study records sporophytes from the current growth season with capsules fully grown, but still green and with lids intact, in early March. Maturity and spore release therefore likely occur during spring or summer in Wales. Nyholm (1998) states sporophyte maturity as early summer in Fennoscandia, whilst Griffin (2014) states June—September in Canada. Sporophytes are described as occurring commonly in *B. halleriana*, both in Britain (Hill et al. 2007; Smith 2004) and elsewhere (Nyholm 1998; Hallingbäck et al. 2008), and there were very few non-sporophytic colonies amongst 15 populations surveyed in Canada (COSEWIC 2011). Within the study site, sporophytes were scarce, with only eight counted from the previous season and four from the current season across ten relevés. Amongst bryophytes, spore size of *B. halleriana* is about average, with a diameter of 18– 23 µm (Ceter et al. 2018), and once airborne would be capable of long-distance dispersal by wind, potentially inter-continental (Wilkinson et al. 2011). There are no estimates of spore output per capsule for *B. halleriana*, but there are estimates of 37,500–121,000 for *Bartramia ithyphylla* subsp. *patens* (Brid.) Fransén (Convey 1994; Convey and Smith 1993). Thus, whilst sporophyte production of *B. halleriana* is scarce within the study area, many millions of spores are likely released each season, which could provide sufficient dispersal for, at least, local population maintenance. Indeed, Hedenäs and Bisang (2019) suggest that widespread long-lived perennial mosses with specialised habitat requirements, such as *B. halleriana*, can maintain genetic diversity and persist in the long term, even when sporophyte production is rare.

Results of the present survey, with a count of 143 occupied trees and logs, show the study site supports an exceptional population of *C. hellerianus*, at least in the context of Wales. Bryophyte survey information from other sites in Wales indicate populations of this scarce liverwort are usually small, with less than ten occupied trees or logs typically detected in sites where it has been found (pers. obs.). A notable exception is Coedydd Nedd a Mellte, Brecknockshire (v.-c. 42) and Glamorganshire (v.-c. 41), where recent surveys have documented 81 occupied trees and logs (Motley and Bosanquet 2017), though spread across a much larger area than the present study site. Curiously, *C. hellerianus* was not found within the study site by Newton (1997, 2002, 2008), but was likely over-looked, since there are records from 1967 and 2011. Within old-growth spruce forests in the core range of the species, *C. hellerianus* can attain much higher population density than found presently. For example, in a plot of 0.25 ha in southern Finland, Laaka-Lindberg et al. (2005) report 47 occupied logs, mostly *Picea abies*. However, reported densities are much lower within mature boreal woodland that is managed for timber production (Gustafsson et al. 2004).

Due to availability of deadwood, *C. hellerianus* is considered a typical member of the epixylic bryophyte flora of old-growth boreal conifer forest (Andersson and Hytteborn 1991; Soderström 1988a, 1988b, 1989). Similarly, the sharp contrast in abundance of this species between Allt Rhyd Groes and Allt Penrhiw-iar is clearly associated with availability of large rotten logs. In the latter woodland compartment, where the liverwort occurs at low density, there are frequent long pieces of decorticated rotten logs, but they are generally slender and <20 cm diameter at base. Such logs are occupied very infrequently by *C. hellerianus* in the study area. In Allt Rhyd Groes, rotten logs are larger and more frequent, and population density of the liverwort is correspondingly much higher. It seems the additional 100 yr of deadwood production in Allt Rhyd Groes mostly explains its superior habitat quality for *C. hellerianus*, and that good oak woodland for this species in Wales takes ca. 200 yr to develop from clear-fell. Whilst Laaka-Lindberg et al. (2005) found frequent occurrence of the liverwort on both large and small deadwood in boreal conifer forest, Soderström (1988a) also documents a preference for large rotten logs.

Besides decorticated rotten logs, which is the principal habitat of *C. hellerianus*, a significant portion of colonies within the study site occur on the bark of living trees, mainly *Q. petraea* but also *B. pubescens. Quercus* has been recorded frequently as a phorophyte in Britain (e.g. Bates 2015; Bosanquet et al. 2005; Hill 1988; Motley and Bosanquet 2017; Paton 1999; Woods 2006), but the liverwort also occurs rarely on *Juniperus communis* (Paton 1999) and there is one previous report from *B. pubescens* (Callaghan 2018). In North America, it has been reported as an epiphyte on *Betula alleghaniensis* Britt., *Pinus strobus* L., *Rhododendron maximum* L. and, frequently in some locations, *Tsuga canadensis* (L.) Carrière (Schuster 1969). Of note, *Tsuga heterophylla* has been planted commonly for timber production in Britain, and there are many mature plantations within the range of *C. hellerianus*, but as yet the liverwort has not been found as an epiphyte on this tree.

No perianths or sporophytes of *C. hellerianus* were found during the present study. Indeed, sporophytes have not been reported in Britain and plants are seldom fertile, though both male and female shoots have been found on the same log (Paton 1999). At other locations in Europe, sporophyte production amongst studied populations is generally rare (Hola et al. 2015; Pohjamo and Laaka-Lindberg 2003; Laaka-Lindberg et al. 2006). As with some populations in Central Europe (Hola et al. 2015), it seems populations in Britain are maintained primarily, perhaps often entirely, by vegetative diaspores and that genetic diversity mainly arises from somatic mutations. Notably, lack of sporophyte production in this species is unlikely to limit dispersal ability within the local landscape, since there is continuous and massive production of foliar gemmae, which due to their small size (10–12 µm) are capable of long-range dispersal by wind (Pohjamo et al. 2006).

This is the first study to have investigated the ecology of *B. halleriana* and *C. hellerianus* in oceanic deciduous woodland. In the context of a warming climate (Jenkins, Murphy, et al. 2009; Jenkins, Perry, et al. 2009), further research would be valuable, given both occur at the edge of their range in Wales, being more typical of colder climates. This could usefully focus on the description and measurement of favourable habitat for each species, for example concerning microclimate and substrate characteristics, and associated physiological tolerances, such as related to light, desiccation and freezing.

## Acknowledgements

Many thanks to Sam Bosanquet (NRW), Heinjo During (Utrecht University) and Oli Pescott (BRC Wallingford) for kind assistance.

## Notes on contributors

Dr Des Callaghan is a field bryologist, researcher and photographer, operating as a consultant under Bryophyte Surveys Ltd, working in Britain and further afield. His research is focused on threatened species, taxonomy and conservation ecology.

Jamie Bevan works for Natural Resources Wales as a Senior Reserves Manager, including responsibility for Allt Rhyd y Groes NNR.

